# Labeling fibrin fibers with microparticle sized beads alter single fibrin fiber lysis, external clot lysis, and produce large fibrin aggregates upon lysis

**DOI:** 10.1101/2021.09.22.461388

**Authors:** Christine Carlisle Helms, Najnin Rimi

## Abstract

**Background:** Fluorescent beads are often used as a tool for visualizing fibrin fibers and can mimic the size of microparticles in the blood. Studies showed microparticles alter the appearance and behavior of whole blood clot systems.

**Objectives:** Here we investigate the effect of beads on fibrin fiber lysis and extensibility to enhance understanding of this common research technique and as a biomimetic system for fibrin-microparticle interaction.

**Methods:** We used fluorescence microscopy, atomic force microscopy (AFM), and scanning electron microscopy (SEM) to quantify changes in lysis, extensibility, and clot structure of fibrin fibers and clots in the presence and absence of beads.

**Results and Conclusions:** Fibrin clot structure and lysis were altered in the presence of beads. Fibrin clots formed with beads had a higher fiber density, smaller fibers, and smaller pores. The rate of lysis for clots was reduced when beads were present. Lysis of bead-labeled individual fibers showed that beads, at concentrations similar to those reported for microparticles in the blood, cause a subset of fibers to resist lysis. In the absence of beads, all fibers lyse. These results demonstrate that beads alter fiber lysis through both a change in fibrin clot structure as well as changes to individual fiber lysis behavior. Additionally, the lysis of clots with beads produced large fibrin aggregates. This data encourages researchers to use careful consideration when labeling fibrin fibers with fluorescent beads and suggests that particles binding fibrin(ogen) in the bloodstream may be an underappreciated mechanism increasing the risk of thrombosis.

## Introduction

Fibrin fibers are a major component of blood clots and an important element in the coagulation system. They form an interconnected network of fibers that incorporate red blood cells, white blood cells, and platelets forming a clot or thrombus. The fibrinolytic system works counter to and alongside the coagulation system. During fibrinolysis, the protease, plasmin, cleaves fibrin producing fibrin degradation products and dissolving blood clots. The coagulation and fibrinolytic systems are highly controlled by many positive and negative feedback loops to maintain homeostasis.

Many disease states show an imbalance between coagulation and fibrinolysis. Accelerated or impaired fibrinolysis is associated with thrombosis, atherosclerosis, diabetes, cirrhosis, and obstetric complications [1]. Altered fibrinolysis often manifests as changes in clot lysis time and these changes show up in multiple medical conditions such as recurrent venous thrombosis and myocardial infarction and may contribute to their morbidity [2].

### Mechanisms of fibrin formation and lysis

Fibrin fibers form when plasma soluble fibrinogen is cleaved by thrombin to form fibrin monomers. Fibrin monomers assemble to form protofibrils which then further aggregate to form fibrin fibers in a mesh network. Factor XIII (FXIII), a transglutaminase, forms additional covalent bonds between the monomers and further strengthens the fibers. Plasmin is formed when plasminogen is cleaved at the Arg561-val562 peptide bond by tissue-type or urokinase-type plasminogen activator (tPA and uPA, respectively). The presence of fibrin localizes and enhances the conversion of plasminogen into plasmin, the primary enzyme responsible for the degradation of a clot.

Plasmin is a serine protease that hydrolyzing the carboxyl side of lysines and arginines [2]. Plasmin(ogen) has kringle domains that bind to lysine residues at over 34 different cleavage sites on fibrin. Electron microscopy work and monoclonal antibody studies show plasminogen binds to fibrin near the end of the molecule, possibly bridging monomers across their distal or D:D interaction [3]. Examination of lysed fibers by electron microscopy and confocal microscopy revealed plasmin degrades fibers laterally leaving blunt ends at the end of fibers [4, 5]. This lateral lysis may be attributed to plasmin “crawling” the width of the monomer (∼ 6 nm) to the next adjacent monomer, rather than along the length of the monomer (∼ 23 nm to the next D:D region in the fiber) during lysis. The “crawling” mechanism proposed by Weisel et al included a second kringle site on plasmin that binds to an adjacent monomer before the first kringle binding is released, this aids the lateral motion of plasmin through the fiber [4]. However, not all lateral transections may be complete, Collet et al observed incomplete lateral transection in thick fibers and thick fibers often had an increase in cross-sectional area before lysis completed [5].

### Lysis studies

Studies of fibrinolysis using whole clots have provided insight into the influence of clot architecture on fibrinolysis. The velocity of the lysis front in clots made of tightly packed, thin fibers is slower compared to less dense, thick fibered clots. This is attributed to the ability of tPA and plasmin to permeate less dense clots more easily [5, 6]. Conversely, at an individual fiber level, the rate of fibrin lysis is greater for thin fibers than for thick fibers [5].

Individual fibers are not restricted to the same lysis trends of whole clots since architectural contexts that influence the behavior of the whole clots do not influence individual fibers. Understanding individual fiber behavior allows one to develop better fibrin clot and lysis models used to progress thrombectomy research. It also may elucidate new information and reveal mechanisms involved in lysis and promote drug discovery. For these reasons, efforts have been made to study the lysis of individual fibers [7, 8].

Previous single fiber lysis showed interesting results. Approximately 30 % or more of the fibers underwent elongation during lysis and did not cleave in two [7, 8]. These results displayed a novel lysis pathway and showed fibrin fibers have an inherent tension in the absence of an external strain. The addition of external strain to individual fibers during lysis displayed an inverse relationship between the elongation and strain [8].

These individual fiber lysis studies were performed with fibers labeled externally with beads. Many studies beyond lysis studies have labeled fibrin with fluorescent beads as a technique for fibrin visualization [9–13]. We are interested in determining the effect that external labeling and particles binding to fibrin have on fiber lysis and other clot properties. In addition to gaining insight into the effect the experimental procedure of labeling fibrin with beads may have on the data, labeling of fibrin with beads may mimic the natural interaction of fibrin with microparticles in the blood.

### Microparticles

Microparticles, in the blood, are a physiological example of materials that bind to fibrin and alter fibrin clot structure [14, 15]. Microparticles, or microvesicles, are extracellular vesicles ranging in size from 100 to 1,000 nm. Microparticles have multiple cellular origins, such as platelets, red blood cells, endothelial cells, white blood cells, and potentially tissues where particles are then collected by blood circulation [16]. There is a wide range of reported values for the concentration of microparticles in the bloodstream. Microparticles are heterogeneous and their size distribution straddles the detection limit of quick counting techniques such a flow cytometry. Slower more tedious techniques such as atomic force microscope, electron microscope, and nanoparticle tracking have been used to count microparticles [17, 18]. However, no matter the technique additional concerns arise for the potential of microparticles production or loss during sample processing [18]. Nevertheless, reported values of microparticles in healthy blood range from 200 to over 10^7^ per ul plasma [19–21] with an average diameter of around 100 to 200 nm [19, 20, 22].

Data supports that approximately 60 % of microparticles in healthy donors are derived from platelets [23]. Microparticles derived from platelets have been linked to an increased risk of thrombotic events in patients with immune thrombocytopenia [24]. Their pro-thrombotic nature may be due to any of the following: binding of coagulation factors to their surface, kinetic regulation of thrombin production, low-grade thrombin production, or physical interaction with fibrin fibers [15, 23, 25]. Zubairova et al. attributed reduced lysis of clots formed from platelet-free plasma in the absence of microparticles to the thrombogenic nature of microparticles as well as their physical interaction with fibers [15].

This work studies the implications of physical interactions between fibrin and beads. Labeling of fibrin with beads of similar size to microparticles may provide insight into the fibrin-microparticle interactions in the bloodstream. We investigate the effect that physical interactions between beads and fibers have on fiber lysis and extensibility. This is a first step in understanding how materials, such as microparticles, that bind fibrin can alter individual fiber properties and hemostasis.

## Methods and Materials

### Proteins and Buffers

Fibrin buffer consisting of 0.05M CaCl_2_, 1M NaCl, and 20mM Hepes at pH 7.4 was used for all sample preparation and dilutions unless specified otherwise. Calcium-free fibrin buffer consisted of 1M NaCl and 20mM Hepes at pH 7.4. Lyophilized human fibrinogen, free of plasminogen, fibronectin, and factor XIII, was acquired from American Diagnostica GmbH (Germany) and was resuspended in a 37 C water bath. Lyophilized labeled fibrinogen, Alexa Fluor 488 labeled human fibrinogen, was obtained from ThermoFisher Scientific (Waltham, MA, USA), reconstituted according to instruction. Human alpha-thrombin, human factor XIII, and human plasmin were acquired from Enzyme Research Laboratories (South Bend, IN). All proteins were aliquoted and stored at −80 C until use.

### Fibrin Sample Preparation

#### Fibrin samples

We formed samples with final concentrations of 0.125 mg/mL 488 labeled fibrinogen, 0.375 mg/mL unlabeled fibrinogen (total fibrinogen concentration of 0.5 mg/ml fibrinogen), 55 Lowey U/ml FXIII, 0.1 U/ml thrombin. The mixture was then transferred to the desired substrate and placed into a humidified chamber in the dark to polymerize for a minimum of 1.5 hours and a maximum of 6 hours.

#### Beads

In samples labeled with beads after polymerization, 20 nm or 200 nm carboxylate-modified fluorescent beads (FluoSpheres, Invitrogen, ThermoFisher, USA) were diluted in calcium-free fibrin buffer. The polymerized sample was rinsed 3 times with calcium-free fibrin buffer, the excess buffer was removed and the diluted bead mixture was added to the sample in the dark at room temperature for 5 minutes. After 5 minutes the sample was rinsed and stored in calcium-free fibrin buffer for data collection. In samples where beads were included during polymerization, calcium-free buffer replace the fibrin buffer in both samples and controls. 200 nm carboxylate-modified fluorescent beads (FluoSpheres, Invitrogen, ThermoFisher, USA) were diluted in calcium-free buffer and added to the fibrin mixture prior to the addition of thrombin to obtain the final concentration indicated.

### Single fiber lysis

Structured surfaces were prepared using a PDMS stamp (Polydimethylsiloxane: Sylgard 184; Dow Corning Corp., Midland MI) formed from a silicon master. The PDMS stamp was pressed into a drop of Norland Optical Adhesive - 81 (Norland Products, NJ) on the top of a microscope slide, and a ring of optical adhesive was created around the stamp to form a well. The glue was cured under UV light. The stamp was then removed leaving an optical glue substrate composed of transparent ridges 12 µm wide and 10 µm high separated by 12 µm wide trenches on glass coverslips.

We prepared samples as indicated above. After polymerization samples were rinsed thoroughly in calcium-free buffer to remove excess fibrin and leave a single layer of fibrin on the surface of the substrate. We then labeled the samples with beads, rinsed them, and stored them in buffer until lysis. We placed the sample on an inverted fluorescent microscope (Zeiss Axio-vert, Germany), a pre-plasmin image was acquired then excess buffer was wicked away from the sample, and plasmin was immediately added at a final concentration of 0.75 U/ml. Images were taken every 5 minutes for 25 minutes.

### Single fiber extensibility

We formed substrates and samples following the same protocol as used for single fiber lysis measurements. Once samples were formed and labeled with beads, we used a combined inverted fluorescent and atomic force microscope (AFM Asylum Research MFP-3D-Bio) to perform fiber manipulations, as done previously [9, 10, 12, 13]. We extended the fibers by pushing the AFM tip laterally into the center of the fiber using an in-house script that allows precise position control of the AFM tip and records tip position and lateral deflection of the cantilever. In addition, fluorescent movies of fiber manipulations were also recorded. Using AFM tip position data as well as fluorescent microscopy images, we calculated engineering strain, 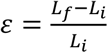, where *L*_*f*_ is the final length of the fiber and *L*_*i*_ is the initial length.

### Clot architecture by SEM

We coated small glass slides with Sigmacote (SigmaAldrich) to reduce surface clot interactions and prepared fibrin samples with beads on the coated slides as indicated above. After polymerization, we took the samples through a series of ethanol dilutions and critical point drying (Tousimis Samdri-795). We then sputter-coated the samples using a palladium target (Denton Vacuum III/IV Desktop). We acquired SEM images using a JEOL SEM at 15 kV, high vacuum, and a magnification of 15,000x.

### External clot lysis

We coated two # 1.5 glass slides with Sigmacote (SigmaAldrich) to reduce surface clot interactions and prepared fibrin samples with beads on one of the coated slides as indicated above. After polymerization, a triple layer of double-sided tape was added to two edges of the second glass slide and the slide was placed on top of the sample, sealing the sample between the slides. We placed the sample on an inverted fluorescent microscope and located the leading edge of the clot. Plasmin was diluted to 1.5 U/ml and 10 ul was added to the edge of the clot using a gel loading pipette tip. After the addition of plasmin, a time point 0 measurement was immediately taken, followed by images every minute until the leading edge of the clot was no longer in the field of view of the microscope.

## Results

### Single fiber elongation during lysis is an artifact of sample preparation

To our knowledge, all previous studies examining single fiber lysis visualize fibrin fibers by topically labeling them with carboxylate modified 20 nm fluorescent beads after fiber polymerization. Following previous protocols, when we labeled fibers with 4.55*10^8^ beads per ul, a 10,000-fold dilution, we obtained similar results to these studies. When exposed to plasmin, the fibers labeled with beads either lysed, redistributed their tension, or showed no change (Figure 1).

**Figure 1.**
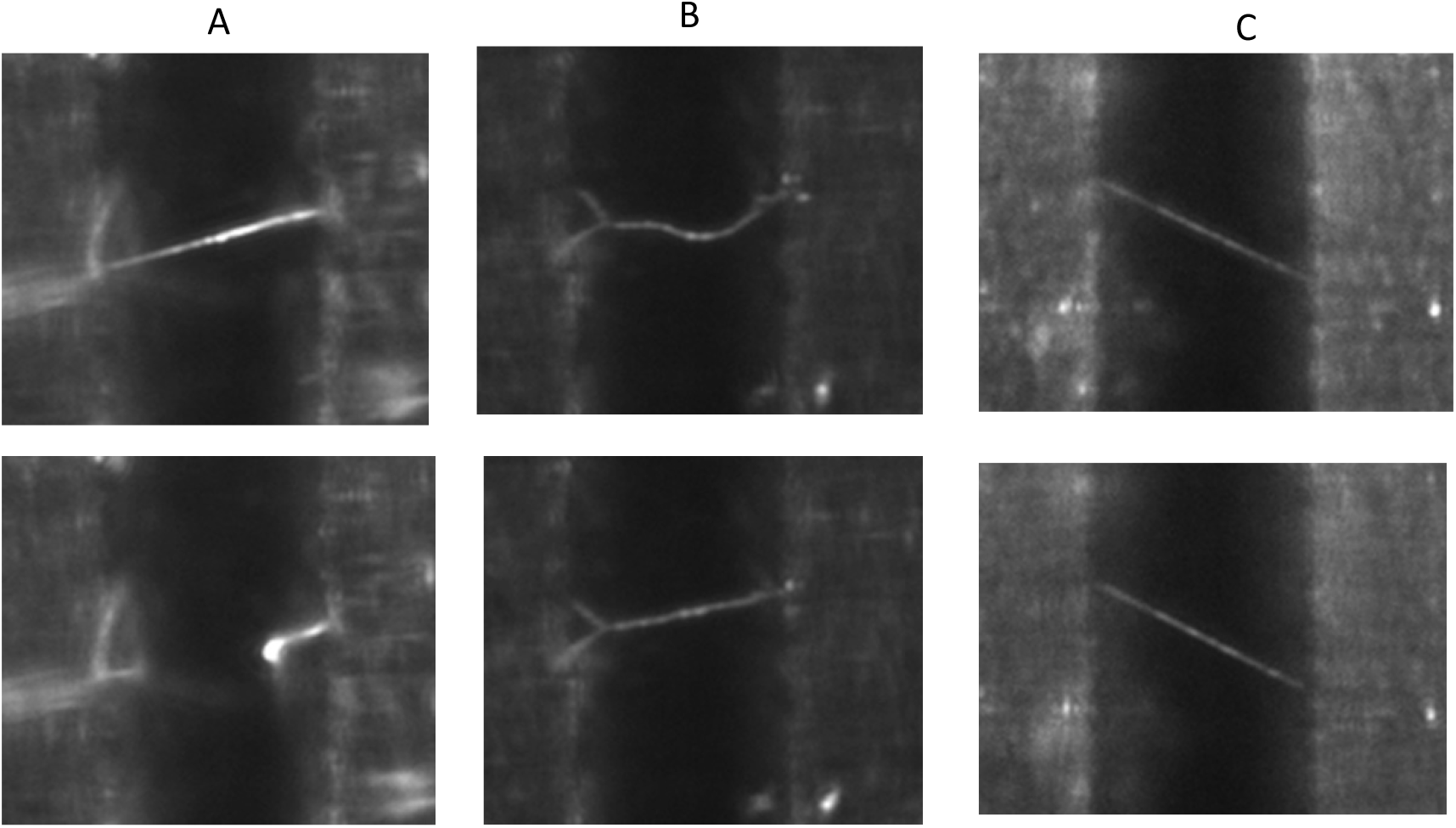
Fluorescent microscopy images of fibers formed from 25 % 488 conjugated fibrinogen and labeled with 20 nm beads at a concentration of 4.55*10^8^ beads per ul. Images show before and after pictures of fibers that (A) lysed, (B) redistributed tension, and (C) showed no change.

Approximately 60 % of individual fibers lysed in two. Among, the 40 % that did not lyse, 18% redistributed their tension and 22 % showed no change (Figure 2). Here our results differed slightly from those reported, as previous work saw all fibers lyse or redistribute at the plasmin levels in use [7]. In addition, fibers that redistributed sometimes elongated but other times they remained taught but changed orientation. We defined redistribution of tension as any change in fiber shape (see Figure 1B). We believe that fibers that changed their orientation yet remain taught, may have done so due to branching along the length of the fiber that was not visible in the plain of view of the microscope.

**Figure 2.**
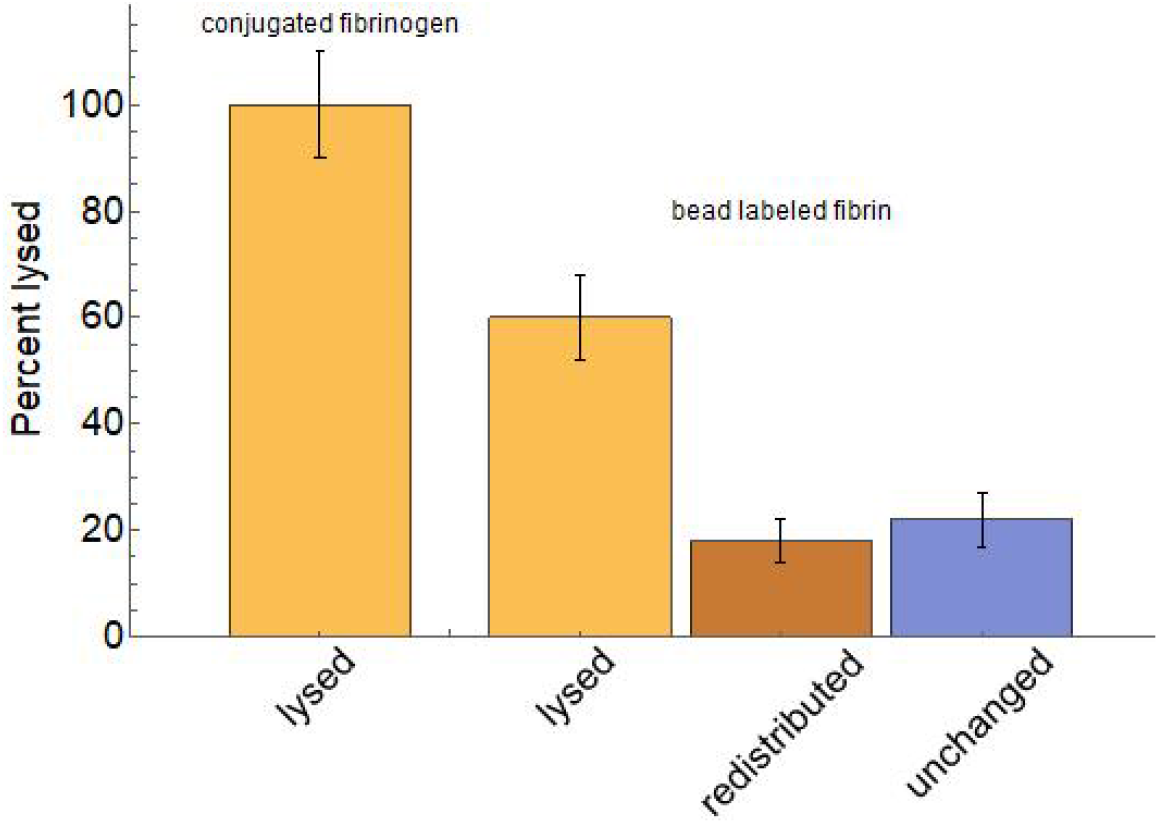
Labeling fibrin fibers after formation with 4.55*10^8^ beads per ul caused some fibers to redistribute their tension or remain unchanged instead of lysing in the presence of 0.75 u/ml of plasmin. The percentage of fibers lysing, redistributing tension, and remaining unchanged are shown in the graph above. (minimum of n = 40)

When we studied the lysis of individual fibrin fibers that were not labeled by beads, all fibers cleave in two typically within 10 minutes of the addition of 0.75 u/ml of plasmin. This suggests that redistribution of tension and the lack of change in fibers upon the addition of plasmin was a result of the external labeling of fibrin fibers with beads.

Indeed, we saw a redistribution of tension in a small percentage of the fibers labeled with beads. This data was consistent with the previously reported elongation of individual fibers during lysis with plasmin [7, 8]. It suggests that in some situations the beads may act to bridge two parts of a lysed fiber together.

### Inhibition of lysis is dependent on bead size and concentration

We compared the effects of 20 nm and 200 nm diameters beads on fiber lysis. We selected the 20 nm beads to reproduce and expand on data from previous single fiber lysis studies. Data compiled by Chandler showed a wide distribution of microvesicles diameters with the maximum of the diameter distribution occurring around 200 nm [22]. Therefore, we choose the 200 nm beads to mimic the peak diameter of microvesicles in the bloodstream.

The previously used 10,000x dilution of 20 nm beads to label fibrin fibers results in a concentration of 4.55*10^8^ beads per ul. As shown in figure 1, this concentration of 20 nm beads prevents 37.5 % of fibers from lysing. As we change the concentration of beads the percent of fibers that lyse changes, figure 3A. At the higher bead concentration fewer fibers lysed and most remained unchanged, while at the lowest bead concentration all fibers lysed. At intermediate bead concentrations, we saw the largest percentage (around 20 %) of fibers redistributing their tension.

**Figure 3.**
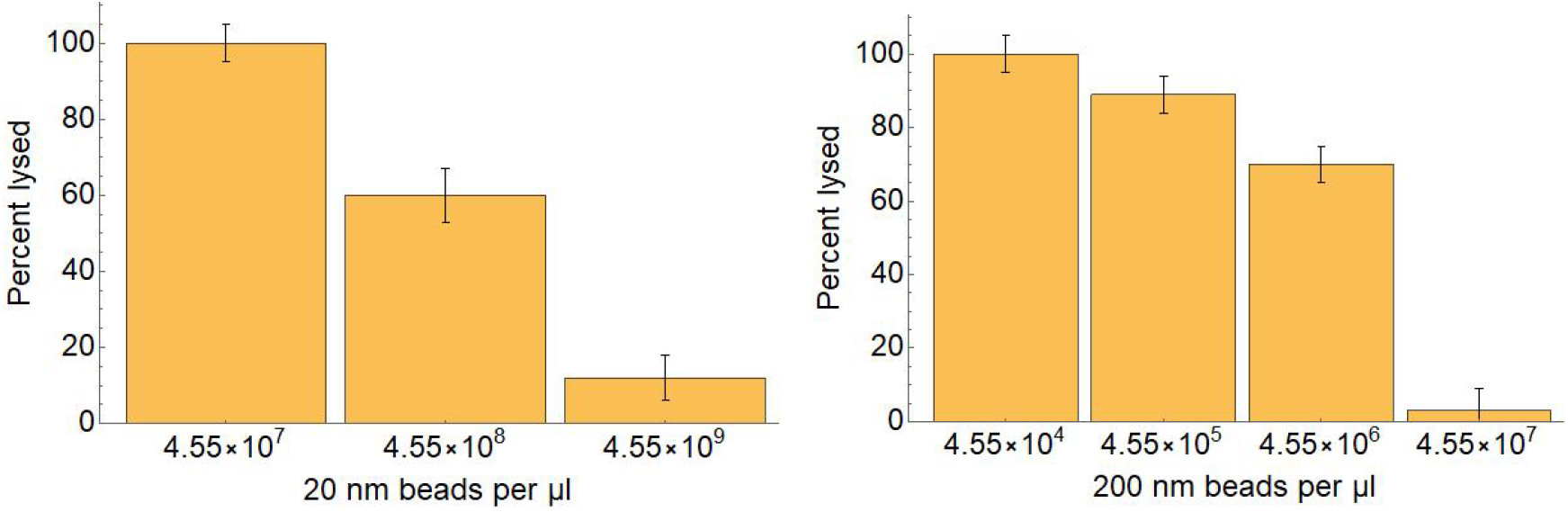
Percent of individual fibrin fibers lysed 25 minutes after the addition of 0.75 u/ml plasmin verse bead concentration. (A) Fibrin fibers are labeled with 20 nm beads. (B) Fibrin fibers are labeled with 200 nm beads. Error bars represent standard deviation based on Poisson statistics.

The concentration of 20 nm beads needed to disrupt fibrin lysis is of the order of magnitude 10^8^ and is above the physiological concentration of microvesicles reported in the bloodstream. However, when we used 200 nm beads to label fibrin, concentrations of beads at and below those reported for microvesicles in the bloodstream were sufficient to disrupt individual fiber lysis, figure 3B. 200 nm beads began to disrupt fibrin fiber lysis at concentrations of 10^5^ beads/ul.

### Beads do not alter fibrin extensibility

As suggested above, interactions between beads and fibers may provide mechanical support to fibers and help hold fibers together during lysis. In other words, the rearrangement of tension in the fibers during lysis suggests the beads may act like a patch placed over a tear. If beads behave in this manner they could reinforce fibers and alter fiber their mechanics under strain. We measured the strain at rupture, or extensibility, of fibers with and without beads to determine if beads alter single fiber extensibility. Using various concentrations of 200 nm beads, we found no significant difference in the extensibility of fibrin fibers labeled with beads to those without beads (Table 1).

**Table 1.** The average extensibility, or strain at rupture, and standard deviation for fibrin fibers labeled with various concentrations of 200 nm beads. Extensibility was defined as the engineering strain at which the fiber ruptured.

| Extensibility data |  |
| --- | --- |
| No beads | 130 % ± 40 % |
| $4.55 \times 10^6$ beads /ul | 130 % ± 60 % |
| $4.55 \times 10^7$ beads /ul | 140 % ± 70 % |

### Beads alter fibrin clot structure

In the data presented above, fibrin polymerized in the absence of beads, and then subsequently beads were added to externally label the fibers. In the following sections, we add beads before fiber polymerization. This resembles physiological conditions where bead-like microvesicles are present in the bloodstream before, during, and after fibrin fiber formation. We formed fibrin clots in the presence and absence of 200 nm beads and found beads altered clot structure.

We defined clot density as the percent of pixels in an SEM image occupied by fibrin fibers. We found no significant difference in clot density between clots made in the presence and absence of beads. However, the average diameter of fibers changed in the presence of beads. When beads were present the average diameter of fibers decreased to 58 nm +/- 15 nm from 74 nm +/- 25 nm, in the absence of beads (p < 0.001, Figure 4). The decrease in fiber diameter, but not in clot density, indicates an increase in the fiber density (or the number of fibers in a region). In other words, if the percent of pixels occupied by fibers remained the same, but the fibers became smaller you would need a larger number of fibers or a greater fiber density to maintain the same number of occupied pixels or clot density. It also indicated a decrease in average pore size in the clot. Therefore, clots formed in the presence of beads were comprised of a larger number of smaller diameter fibers and the clots had smaller pore sizes than those without beads.

**Figure 4.**
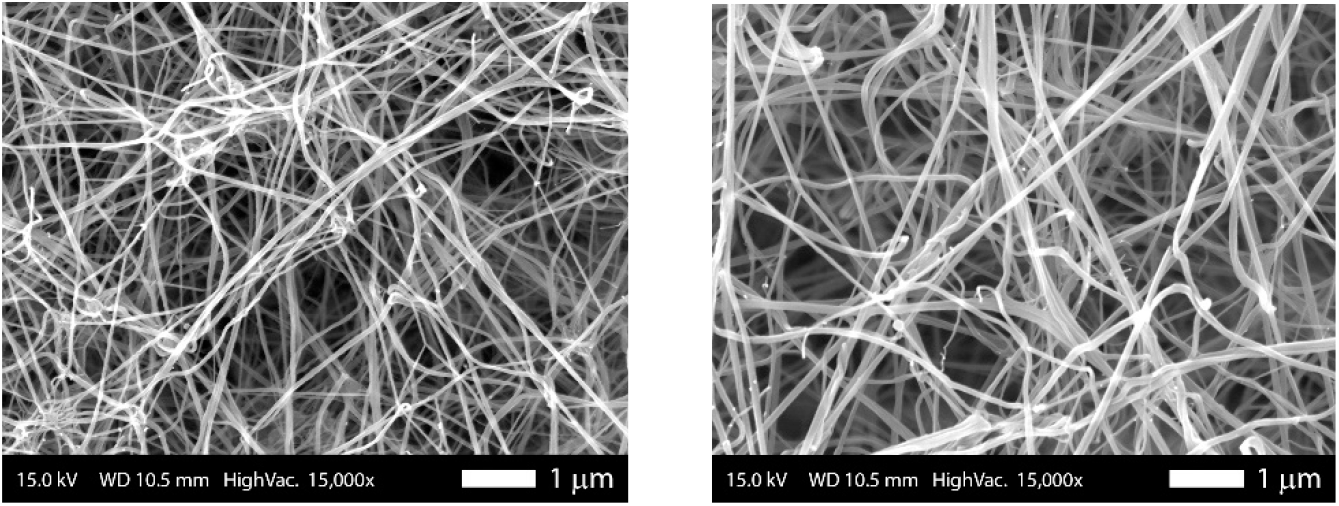
Scanning electron micrographs of fibrin clots formed in the presence (A) and absence (B) of 200 nm beads at a concentration of 4.55*10^6^ beads per microliter. The average clot density was unchanged by the presence of beads. The average fiber diameter decreased in the presence of beads (A) compared to samples without beads (B).

### Beads decrease the rate of clot lysis

We measured external clot lysis using fluorescence microscopy. A location 300 microns from the leading edge of the clot was identified immediately following the addition of plasmin at time point t = 0 min. We recorded the distance of the leading edge of the clot from this defined location every minute until the clot receded beyond the location (Figure 5). Clots formed in the presence of beads lysed slower than clots without beads. The rate of lysis for clots without beads, determined by least-squares fitting, was 73±8*μ*m/min while the rate of lysis for clots with beads was 52±5 *μ*m/min (best fit ± standard error of the mean, n = 8).

**Figure 5.**
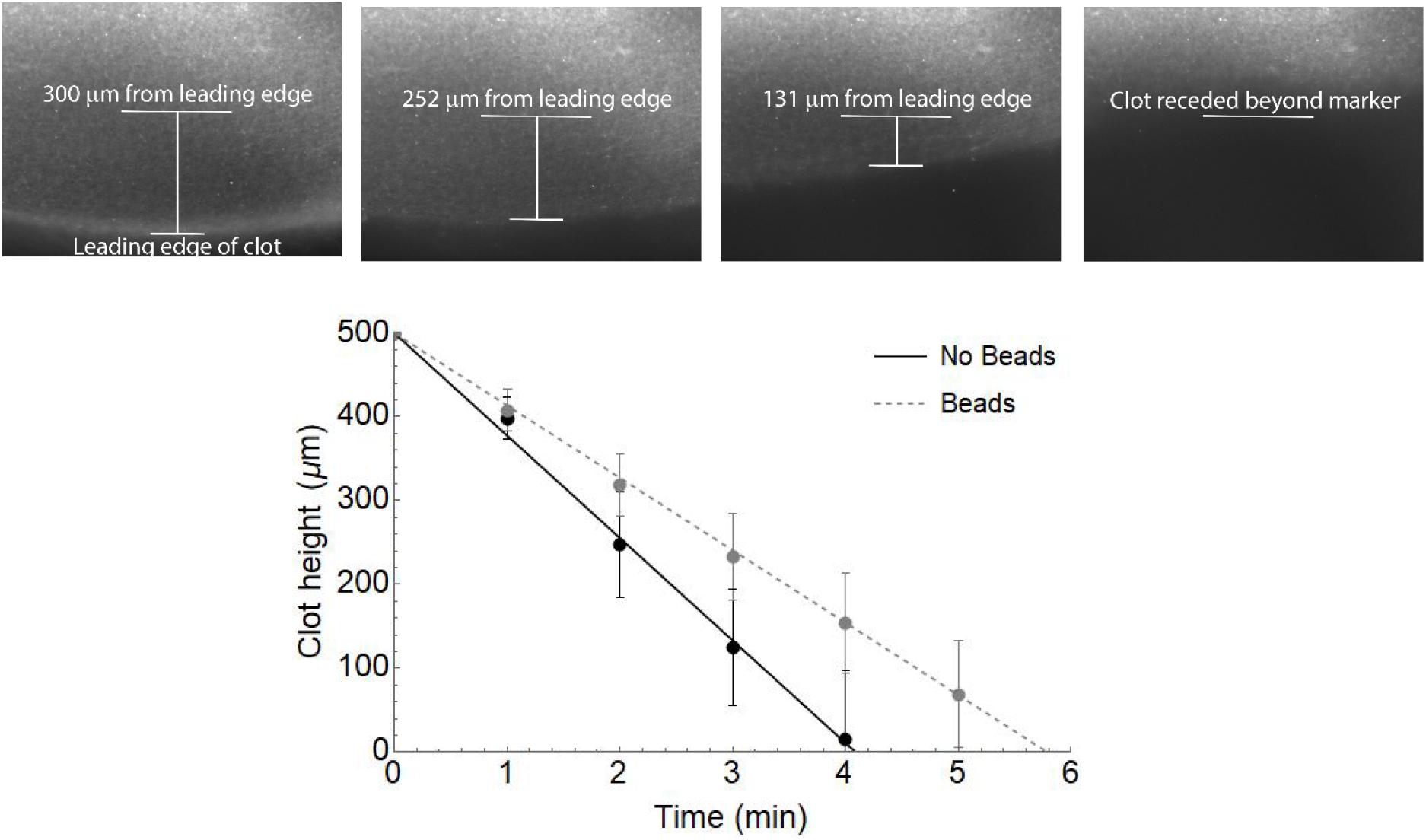
(A) Fluorescence microscopy images of fibrin clot receding assay. A position 300 microns from the leading edge of the clot was defined immediately following the addition of plasmin (long horizontal bar). The distance between the leading edge and this location was recorded every minute. (B) Data from the receding clot assay were plotted versus time. The rate of clot lysis is given by a linear least-squares fit to the data, 73±8*μ*m/min for clots without beads and 52±5 *μ*m/min for clots with 200 nm beads at a concentration of 4.55*10^6^ beads per ul during clot formation. Circles represent an average of eight data points and error bars give the standard error of the mean.

In addition to an altered rate of lysis, clots with beads formed large fibrin aggregates during lysis (Figure 6). Aggerates like the ones pictured in Figure 6 were seen in all external clot lysis assays that included beads. A few aggregates were followed for up to 30 minutes post plasmin. During this time the clot receded however, once the aggregates formed they did not reduce in size. Similarly, fibrin aggregates were not seen during the lysis of clots without beads.

**Figure 6.**
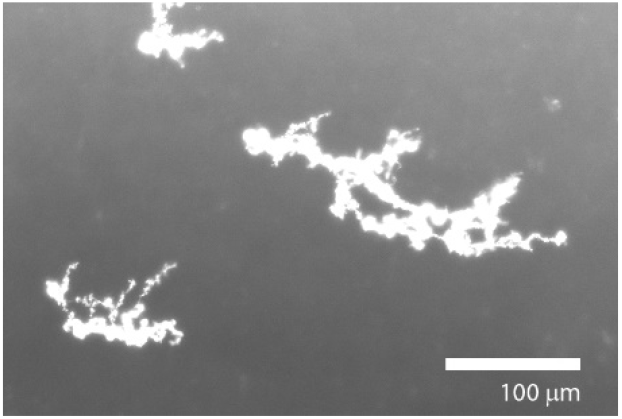
Fluorescence microscopy image of fibrin aggregates formed during the lysis of fibrin clots containing beads.

## Discussion and Conclusion

Our data show that previously reported elongation of individual fibrin fibers during lysis is a result of labeling fibrin fibers with fluorescent beads for visualization. Upon more detailed investigation elongation or redistribution of tension within the fiber occurs only at intermediate concentrations of beads. When fibers are labeled with a high concentration of beads the fibers show no change in structure upon the addition of plasmin. At low concentrations of beads, all fibers lyse upon the addition of plasmin. The concentration of beads needed to elicit a particular behavior, in other words, the concentrations that define low, intermediate, and high, depends on the diameter of the beads. Larger beads affect fibrin lysis at lower concentrations than smaller beads.

20 nm diameter FluoSpheres have frequently been used to label individual fibrin fibers during research [7, 9, 11, 13, 26–28]. These studies do not consider the effect of fluorescent labeling on their results. Our lysis data suggest that labeling of fibrin with beads affects the behavior of fibrin and therefore, studies and data using this labeling technique must be interpreted accordingly. In addition to affecting the interpretation of previous data, the results of this work suggest an importance of the binding of particles to fibrin. In particular, the beads used in many experiments in this study were selected to mimic the peak size of microparticles in the bloodstream. Studies on the size distribution of microparticles indicate a wide range of diameters with a peak diameter around 200 nm [19, 20, 22]. Therefore, we used 200 nm beads to label fibrin, unless otherwise indicated.

Beads with a diameter of 200 nm began to alter single fiber lysis at concentrations of 10^5^ beads/*μ*l. This concentration is within the reported physiological range of microparticles in the blood [19–21]. At this and higher concentrations, the labeling of fibrin fibers with beads prevented fiber lysis, with fibers showing either no response to plasmin or redistribution of tension upon the addition of plasmin.

We propose two possible mechanisms for the prevention of fiber lysis in the presence of beads. First, the beads may block the plasmin binding sight on fibrin and sterically hinder the binding of plasmin to the fibrin fibers, therefore, preventing lysis. Second, plasmin may cleave the fibrin fiber in two but the beads, coated with pendant carboxylic acids, may span the cleavage site and bind to the two halves of the fiber holding the lysed segments together. If the beads were sterically preventing plasmin from binding fibrin fibers, then one would expect fibers to remain unchanged when exposed to plasmin. However, if beads were working to hold the fibers together then redistribution of tension may occur as beads become the sole support for the tension when fibrin is cleavage.

The redistribution of tension we see in a subset of bead-labeled fibers suggests beads may be able to bear tension along the length of the fiber. In this same manner, bead-labeled fibers under strain could use the beads to provide additional mechanical support. Therefore, we tested the effect of labeling fibrin with beads on fiber extensibility. Surprisingly, labeling fibers with beads did not affect maximum fibrin fiber extensibility. This finding validates the results from previous extensibility studies and suggests that beads may not significantly affect the results of other fiber mechanical measurements such as modulus.

The effect of beads on the lysis of fibrin clots goes beyond alterations to individual fiber lysis. In measurements of external clot lysis, the presence of beads altered the rate of lysis. Along these lines, Zubairova et al. formed clots from platelet-free plasma and filtered platelet-free plasma that removed microparticles and showed plasma with microparticles lysed slower than plasma without microparticles [15]. Our data agree with these findings and the agreement supports the idea that beads may mimic microparticles behavior. Alternately, previous studies using much smaller beads, 5 nm, showed no effect of these beads on clot properties [29]. Our data may explain these seemingly different results, as we found a higher concentration of smaller beads is needed to alter fibrin lysis behavior.

We proposed two potential mechanisms for the decreased rate of fibrinolysis in the samples with beads. First, the decreased rate of lysis may be due to the structural changes to the fibrin clots in the presence of beads. Our SEM data suggests a higher fiber density and smaller pore size in clots formed with beads which may slow the diffusion of plasmin into the clot and therefore decrease the rate of lysis. A reduced rate of clot lysis is commonly seen in clots with this dense structural architecture [29, 30]. Second, the subset of fibers that do not lyse in the presence of beads, shown in our individual fiber data, may affect the diffusion of plasmin through the clot by physically blocking plasmin movement.

Additionally, the subset of fibers that do not lyse may result in lysis-resistant large fibrin aggregates during external clot lysis. The appearance of various-sized fibrin aggregates was evident in samples formed with beads. These aggregates did not diminish in size throughout the 20-minutes they were optically tracked. Similar fibrin aggregates were not found during the lysis of clots without beads. Fibrin aggregates in the blood could promote a prothrombotic state and encourage embolism.

In this work, we showed the effect of labeling fibrin with fluorescent beads on individual fiber lysis. Adding clarity to the interpretation of previous data on individual fiber lysis [7, 8]. Our data shows the novel elongation of fibers seen in previous studies was a result of fiber labeling with beads. The impact of beads shown through this work encourages researchers to consider how labeling fibrin may alter other fiber measurements. Additionally, bead-labeled fibrin may have a physiological parallel in microparticles. Therefore, understanding the changes in clot architecture and lysis due to beads may have great clinical value. Our results suggest that a subset of fibers labeled with beads do not lyse and may add to some of the changes seen during external clot lysis such as reduced lysis rate and fibrin aggregate formation.

## Author Contributions

C.C. Helms performed study concept and design. All authors provided acquisition, analysis, and interpretation of data. All authors contributed to the writing, review, revision and approval of the final manuscript.

## Disclosure of Conflict of Interest

C.C. Helms and N. Rimi declare that they have no conflict of interest.

## Notes

### Competing Interest Statement

The authors have declared no competing interest.

